# Mapping Brain Growth and Sex Differences Across Prenatal to Postnatal Development

**DOI:** 10.1101/2025.03.17.643640

**Authors:** Yumnah T. Khan, Alex Tsompanidis, Marcin A. Radecki, Carrie Allison, Meng-Chuan Lai, Richard A. I. Bethlehem, Simon Baron-Cohen

## Abstract

The perinatal period, encompassing both prenatal and early postnatal stages, is a highly dynamic and foundational phase of brain development. Despite its significance, limited work has tracked brain growth continuously across prenatal to postnatal development. In this study, we analysed one of the largest perinatal MRI datasets from the Developing Human Connectome Project (798 scans from 699 unique individuals: 263 prenatal and 535 neonatal; 380 males and 319 females) to model age-related changes and sex differences in brain volumes from 21 to 45 weeks postconceptional age. We found that total brain volume grew at an increasing rate, with white matter dominating mid-gestational growth and gray matter dominating late-gestational and postnatal growth. Subcortical gray matter structures showed distinct trajectories and earlier peak growth rates compared to cortical gray matter structures. Additionally, sex differences in brain growth patterns were observed, with males showing greater volumetric increases with age compared with females. The findings demonstrate the evolving structural dynamics of perinatal brain development as well as the importance of integrating prenatal and postnatal neuroimaging to map continuous early brain growth trajectories.

## Introduction

The perinatal period, spanning both the prenatal and early postnatal stages, is a highly dynamic and foundational phase of brain development. During this critical window, the brain’s core architecture is established through developmental processes such as cell proliferation, migration, differentiation, synaptogenesis, dendritic growth, and neuronal circuit formation^1,2^. These processes lay the groundwork for lifelong cognitive, behavioural, and health outcomes^3–6^. Despite its significance, this critical window of brain development remains insufficiently studied due to various challenges associated with applying neuroimaging to fetus and infants.

While existing studies have separately mapped prenatal and postnatal brain development^7–11^, it is also informative to track brain growth continuously from prenatal to postnatal development. This approach captures both the continuous nature of perinatal brain development as well as the brain’s response to the prenatal to postnatal transition, enabling the mapping of comprehensive early growth trajectories. While the emergence of large-scale, publicly available datasets has recently enabled investigations into the perinatal development of global brain metrics^12^, an understanding of regional development during this formative period remains limited. It is likely that different regions follow distinct developmental timelines that align with their emerging functional roles, structural organisation, and integration into broader neural networks. For instance, it has previously been suggested that regions responsible for basic sensory functions mature first, followed by associative regions responsible for higher-order cognition^13^. Moreover, growth patterns are likely also impacted by the significant environmental changes between the prenatal and postnatal periods. For instance, genetic, placental and maternal factors (e.g., hormones, nutrition, pregnancy trajectories) are key contributors to prenatal brain development^6,14^. In contrast, postnatal development is further shaped by postnatal life experiences (e.g., sensory stimuli, social interactions) which likely promote growth in specific regions^15^. Studying prenatal and postnatal brain development together can therefore provide insights into how the transition between prenatal to postnatal stages impacts growth dynamics in early development.

Importantly, the perinatal period is also a critical window for the emergence of sex differences in the brain. Given that sex differences in brain structure are well-documented across the lifespan^16^ and are observed as early as the prenatal and neonatal periods^17,18^, sex-differential growth likely begins early in development. A key factor contributing to these differences is the prenatal testosterone surge, during which male fetuses produce approximately 2.5 times more testosterone than female fetuses in the first and second trimesters of pregnancy^19^. This prenatal surge in testosterone, along with sex differences in placental and neuronal gene expression, are understood as key early biological mechanisms initiating sex differences in brain development^20^. There is also growing evidence suggesting that various perinatal exposures—such as pregnancy complications or maternal drug use^21^— impact brain development in sex-specific ways. Given the high plasticity of the perinatal brain, it is likely particularly receptive to these sex-differential biological and environmental factors. Importantly, sex differences are observed in the prevalence and presentation of different neurodevelopmental and psychiatric conditions^22,23^. This may be at least partly attributable to sex differences in early brain development, as variations in brain structure associated with these conditions overlap with sex differences in brain structure^22,24,25^. Given that perinatal brain development is pivotal to shaping lifespan outcomes^6^, these early differential growth trajectories might explain why sex differences are observed in health outcomes.

Despite being a critical stage for brain development, there remains a lack of well-powered research on the perinatal period. Existing studies often do not incorporate both prenatal and postnatal scans when modelling early brain growth, obscuring a full picture of early developmental trajectories. The Developing Human Connectome Project (dHCP) dataset^26^, which includes both prenatal and neonatal MRI scans, offers a unique opportunity to address this research gap. In the present study, we analysed 798 partially longitudinal prenatal and neonatal MRI scans from 699 unique individuals (263 prenatal, 535 neonatal; 380 males, and 319 females) from 21 to 45 weeks post-conception to model age-related structural changes and sex differences in perinatal brain volumes.

## Results

In the present study, we used dHCP prenatal and postnatal scans to map continuous perinatal developmental trajectories of global and regional brain volumes using mixed-effect models. A subset of the sample was partially longitudinal (N=97 out of 699). Within this group, 78 participants had both fetal and neonatal scans, 16 had two fetal scans, one had three fetal scans, and two had two neonatal scans. Two sets of analyses were conducted – one using absolute brain volumes and the one using proportional brain volumes, where the brain volume of interest was expressed as a proportion or fraction of total brain volume. In a pre-analysis step, Bayesian Information Criterion (BIC) values were computed for each global and regional volume (separately for absolute and proportional analyses) to determine whether a linear, quadratic, or cubic model best described its growth trajectory (Supplementary Tables 3 and 4). Mixed-effects models with random intercepts modelled at the subject-level were then fit to each measure using the selected model. The models simultaneously included main effects of age, sex, and sex-by-age interactions. Full model comparison results are reported in Supplementary Materials. Each analysis was FDR-corrected for multiple comparisons using a significance threshold of ≤ 0.05^27^. Standardised beta coefficients were also calculated for each analysis to denote standardised effect sizes, reported fully in the Supplementary Materials.

### Absolute Global Analysis

The growth trajectories of total brain volume (TBV), total white matter volume (WMV), and total gray matter volume (GMV) were best described by quadratic models. Total brain volume (Age: µ = 17995, *SE* = 270, *p_FDR_* <0.001, Age^2^: µ = 179, *SE* = 35, *p_FDR_* <0.001) and total gray matter volume (Age: µ = 12641, *SE* = 156, *p_FDR_* <0.001, Age^2^: µ = 277, *SE* = 20, *p_FDR_* <0.001) both showed increasing growth rates with age, while total white matter volume (Age: µ = 5345, *SE* = 125, *p_FDR_* <0.001, Age^2^: µ = −97, *SE* = 16, *p_FDR_* <0.001) showed decreasing growth rates. All of these volumes showed linear sex-by-age interactions with significantly greater increases in males (TBV: µ = 1765, *SE* = 387, *p_FDR_* <0.001, GMV: µ = 1216, *SE* = 224, *p_FDR_* <0.001, WMV: µ = 556, *SE* = 179, *p_FDR_* = 0.003). All quadratic sex-by-age interaction terms were non-significant (TBV: µ = 35, *SE* = 50, *p_FDR_* = 0.54, GMV: µ = 35, *SE* = 50, *p_FDR_*=0.54, WMV: µ = −9, *SE* = 23, *p_FDR_* = 0.71).

Intracranial volume (ICV) (Age: µ = 16891, *SE* = 422, *p_FDR_* <0.001, Age^2^: µ = 317, *SE* = 96, *p_FDR_* <0.001, Age^3^: µ = 30, *SE* = 7, *p_FDR_* <0.001) and cerebrospinal fluid (CSF) (Age: µ = −1078, *SE* = 175, *p_FDR_* <0.001, Age^2^: µ = 142, *SE* = 37, *p_FDR_* <0.001, Age^3^: µ = 30, *SE* = 3, *p_FDR_* <0.001) both showed cubic trajectories. CSF volumes declined at 30-40 weeks and then continued increasing at an accelerating rate from 40 weeks onwards. ICV showed a significant linear sex-by-age interaction with greater increases in males (µ = 1621, *SE* = 599, *p_FDR_* =0.01), but non-significant quadratic (µ = 243, *SE* = 134, *p_FDR_* = 0.11) and cubic (µ = 15, *SE* = 10, *p_FDR_* =0.18) interaction terms. CSF showed significant quadratic (µ = 142, *SE* = 37, *p_FDR_* <0.001) and cubic (µ = 30, *SE* = 3, *p_FDR_* <0.001) sex-by-age interactions, indicating a more pronounced cubic trajectory in males (see Figure 1).

**Figure 1.**
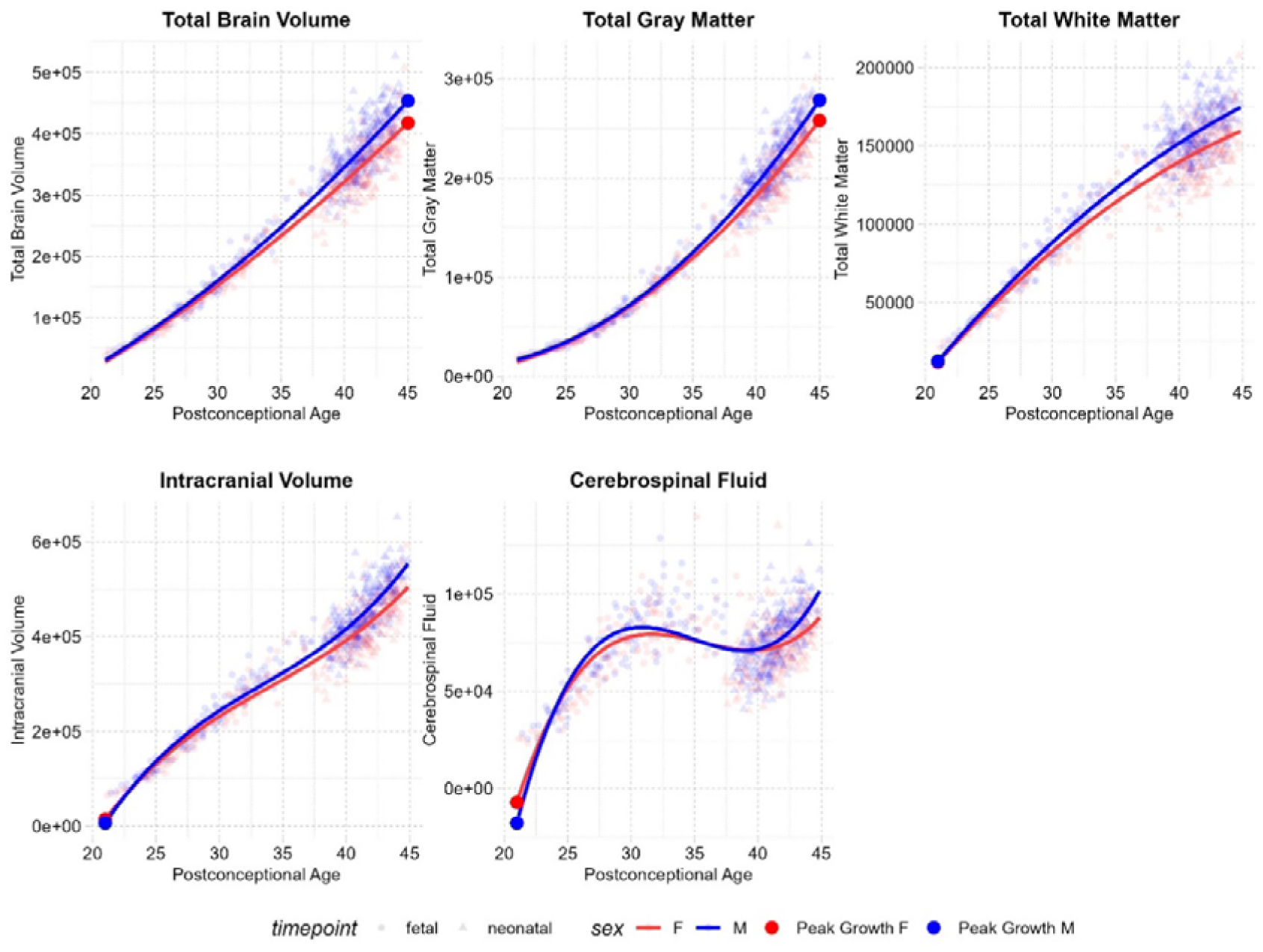
Global Growth Trajectories

### Proportional Global Analysis

In order to understand changes in the relative contributions of gray and white matter to overall brain growth while avoiding model collinearity, total gray and white matter volumes were analysed as proportions of total brain volume (Figure 2). Cubic models best fit these proportional trajectories. Across the studied period, total gray matter proportions increased (Age: µ = 0.01, *SE* = 0.00, *p_FDR_*<0.001, Age^2^: µ = −0.00, *SE* = 0.00, *p_FDR_* = 0.02, Age^3^: µ = −0.00, *SE* = 0.00, *p_FDR_*<0.001) while total white matter proportions decreased (Age: µ = −0.01, *SE* = 0.00, *p_FDR_* <0.001, Age^2^: µ = 0.00, *SE* = 0.00, *p_FDR_* = 0.02, Age^3^: µ = 0.00, *SE* = 0.00, *p_FDR_* <0.001). White matter showed greater contributions (>50%) to total brain volume before 35 weeks, while gray matter showed greater contributions after 35 weeks. All sex-by-age interactions were non-significant for both gray (Sex*Age: µ = 0.00, *SE* = 0.00, *p_FDR_*=0.23, Sex*Age^2^: µ = −0.00, *SE* = 0.00, *p_FDR_* = 0.95, Sex*Age^3^: µ = −0.00, *SE* = 0.00, *p_FDR_*=0.65) and white matter proportions (Sex*Age: µ = −0.00, *SE* = 0.00, *p_FDR_* = 0.24, Sex*Age^2^: µ = 0.00, *SE* = 0.00, *p_FDR_* = 0.95, Sex*Age^3^: µ = 0.00, *SE* = 0.00, *p_FDR_* = 0.65).

**Figure 2.**
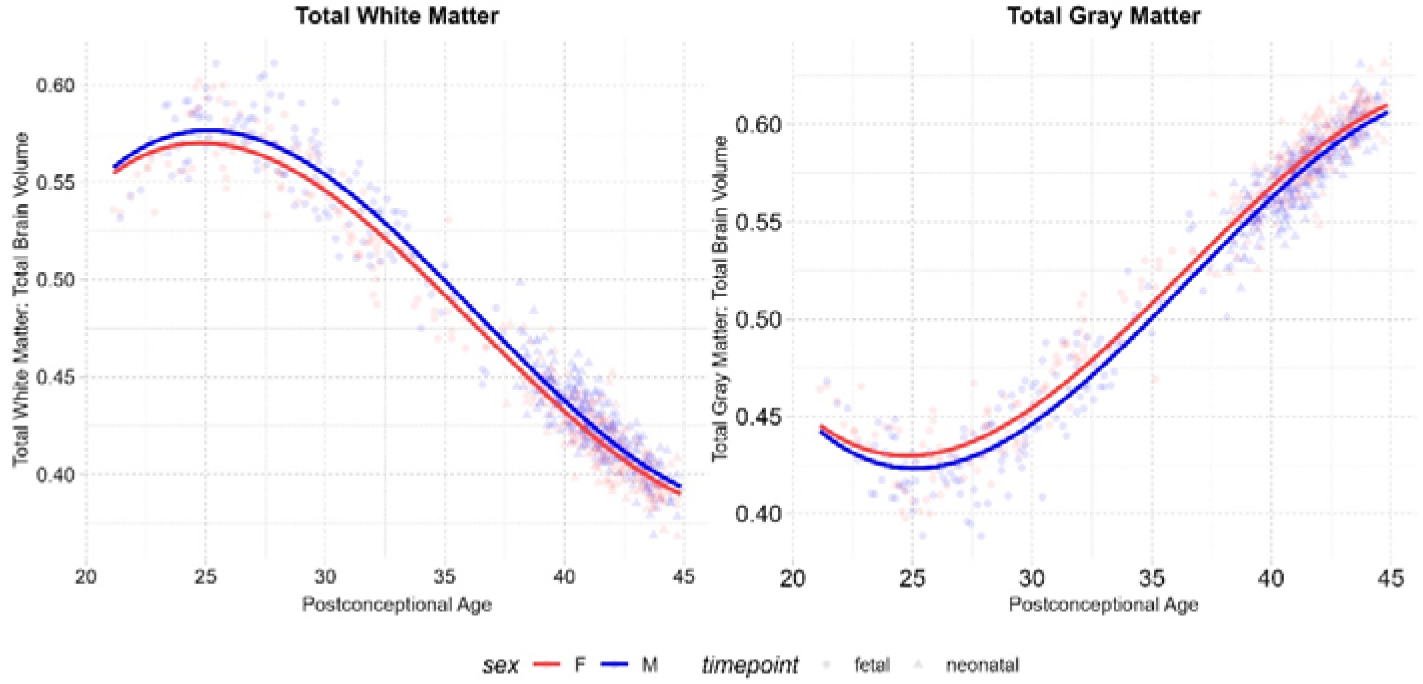
Proportional Gray and White Matter Trajectories

### Absolute Regional Analysis

All regional analyses reported here are on absolute brain volumes. Regional trajectories as a proportion of total brain volume are reported in Supplementary Materials (Supplementary Table 2). Following the global trends reported above, gray matter in the four cortical lobes showed quadratic trajectories with increasing growth rates, while white matter showed quadratic trajectories with decreasing growth rates (Table 1). Results and figures from the full parcellation (87 structures) are reported in Supplementary Materials (Supplementary Table 1). Various subcortical regions, such as the amygdala, thalamus, and basal ganglia structures, showed cubic growth trajectories. These trajectories were characterised by an initial phase of accelerating growth which peaked during the third trimester, followed by a subsequent phase of decelerating growth postnatally. The corpus callosum and cerebellum showed quadratic trajectories with increasing growth rates, while the hippocampus showed linear growth throughout the studied period (Table 1 and Figure 3).

**Figure 3.**
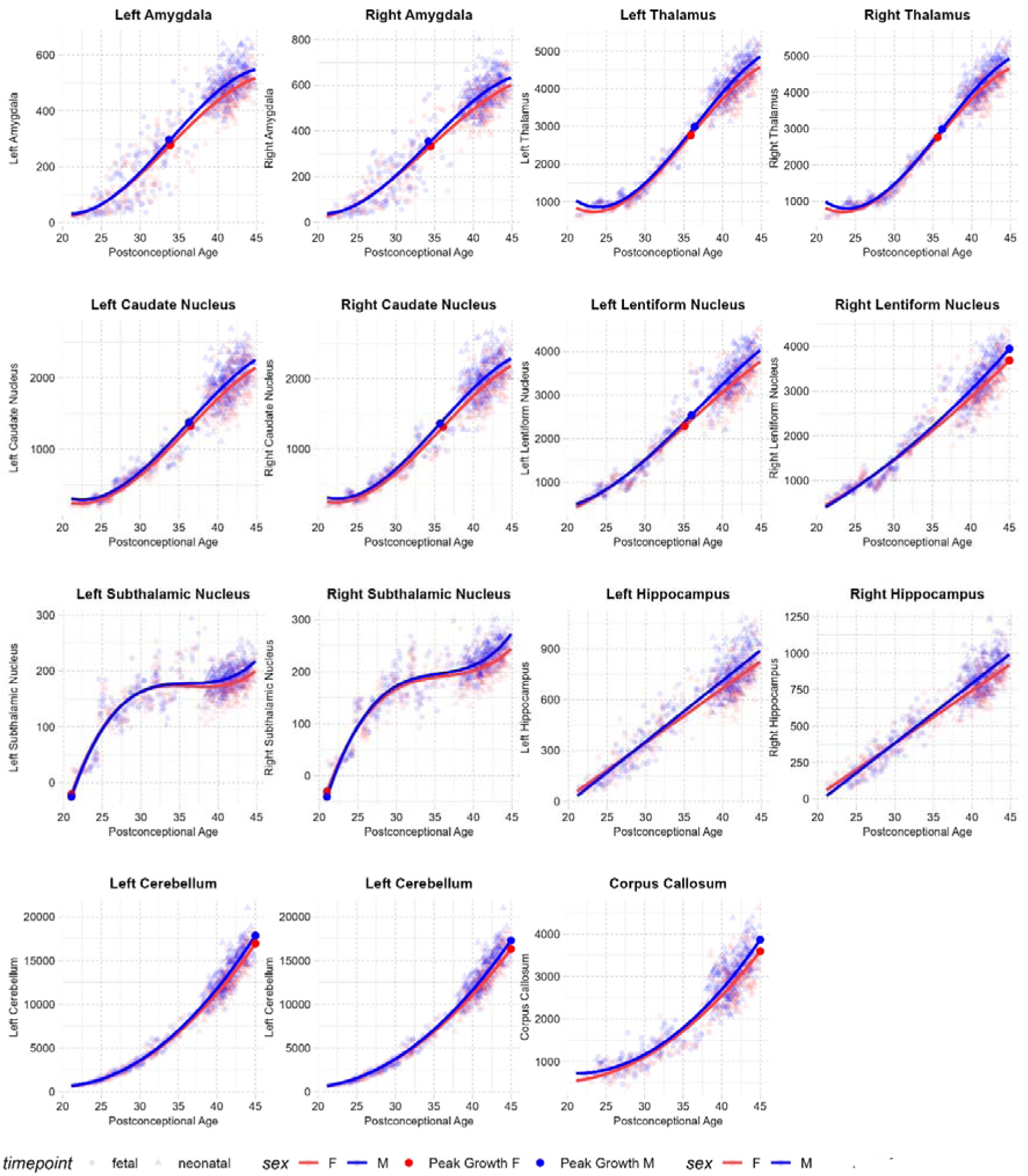
Subcortical Growth Trajectories

**Table 1.**
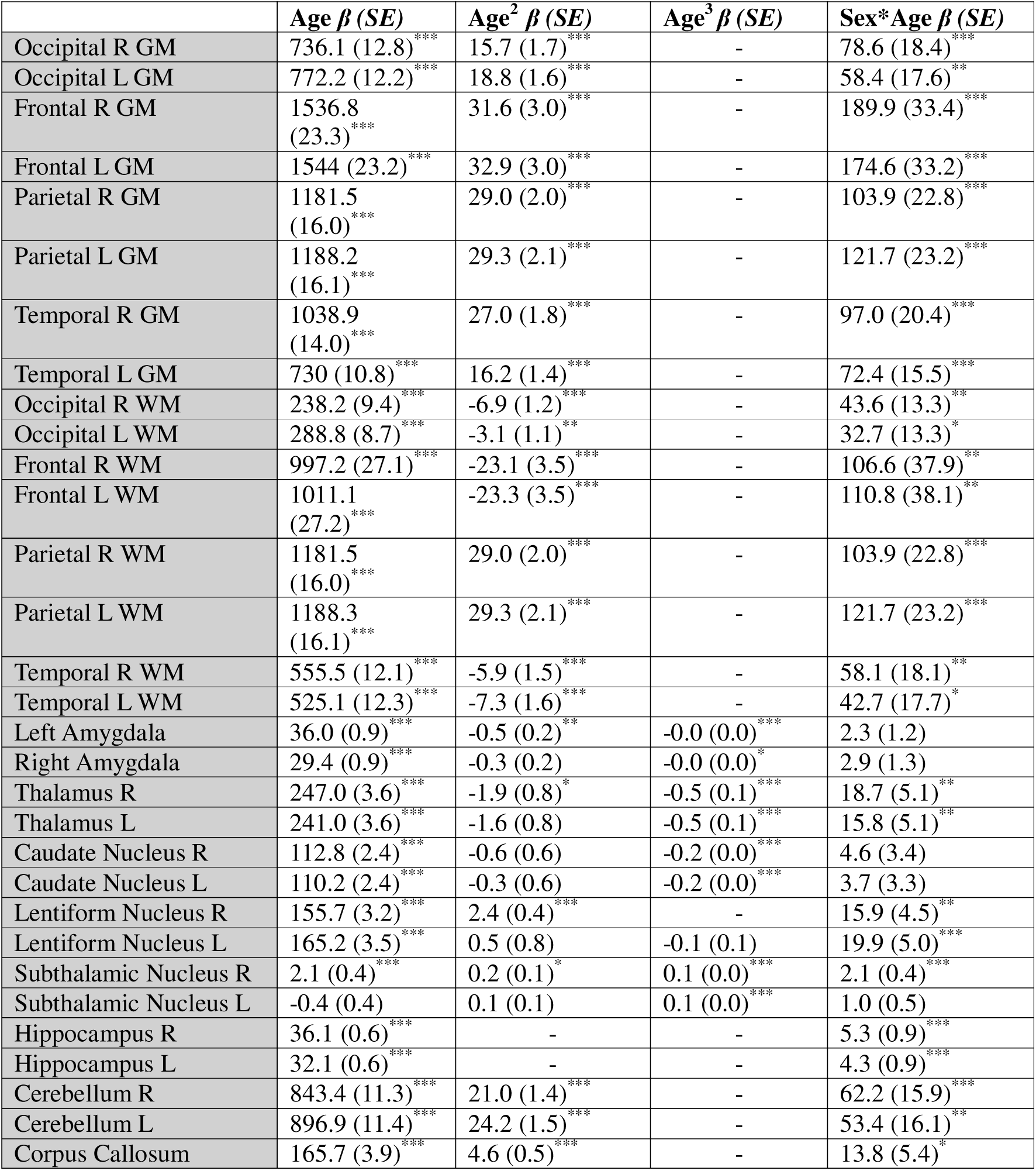
Regional Models. *Mixed-effect model outputs, showing beta coefficients (, standard errors (SE), and significance levels (* p_FDR_<0.05, ** p_FDR_ <0.01, *** p_FDR_ <0.001) of the linear, quadratic, and cubic age terms and linear sex-by-age interactions. In sex-by-age interactions, positive interaction coefficients indicate faster growth in males. All p values are FDR-corrected across the full regional parcellation (87 regions, see Supplementary Materials). R = right hemisphere, L = left hemisphere, GM = Gray Matter, WM = White Matter*.

### Absolute Regional Sex Differences

All sex differences reported here are on absolute brain volumes. Sex differences on regional trajectories as a proportion of total brain volume are reported in Supplementary Materials (Supplementary Table 2). Most regions showed linear sex-by-age interactions with significantly faster growth in males (Table 1). Significant quadratic sex-by-age interactions (Figure 4) were identified in the anterior parts of the right medial and inferior temporal gyri, left parietal lobes, and posterior parts of the bilateral superior temporal gyri, where males showed more rapid and pronounced gray matter increases compared to females. A significant cubic sex-by-age interaction was identified in the left anterior cingulate gyrus, where the plots show a cubic trajectory in males and a linear trajectory in females.

**Table 2.**
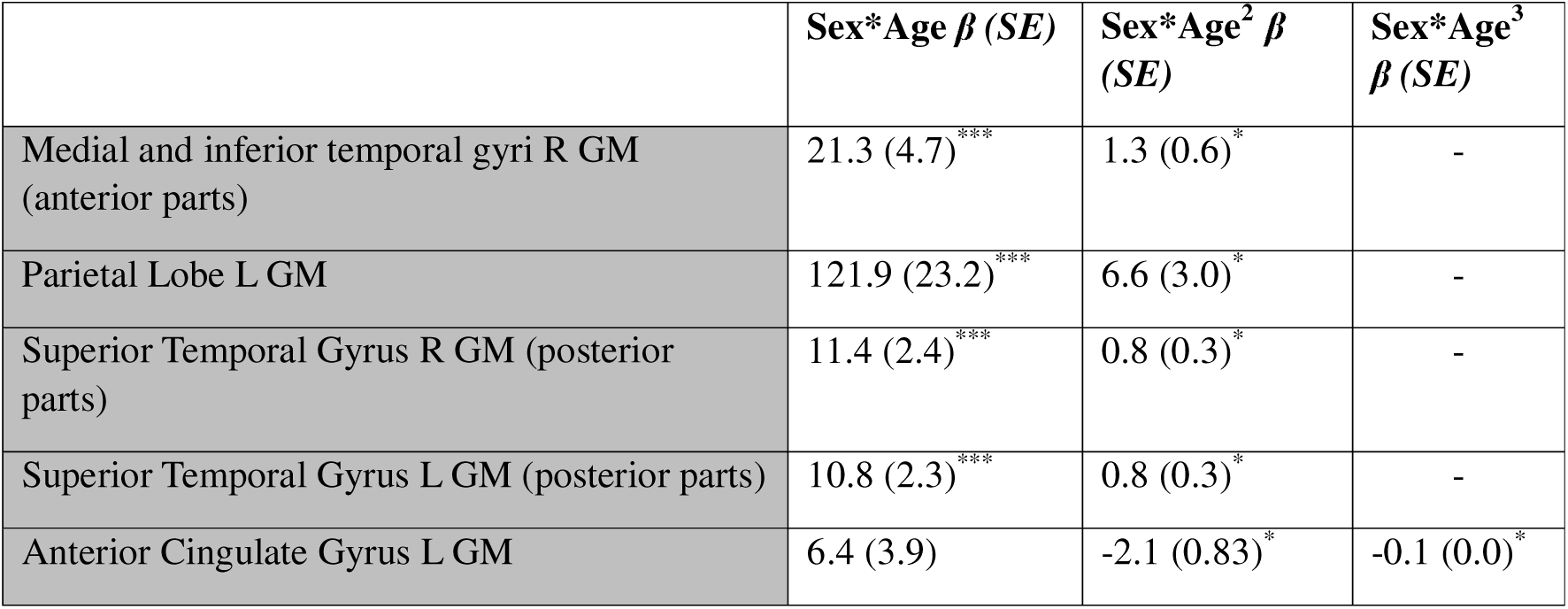
Regions Showing Significant Quadratic and Cubic Sex-by-Age Interactions.

|  | **Sex\*Age $\beta$ (SE)** | **Sex\*Age<sup>2</sup> $\beta$ (SE)** | **Sex\*Age<sup>3</sup> $\beta$ (SE)** |
| --- | --- | --- | --- |
| Medial and inferior temporal gyri R GM (anterior parts) | 21.3 (4.7)<sup>***</sup> | 1.3 (0.6)<sup>*</sup> | - |
| Parietal Lobe L GM | 121.9 (23.2)<sup>***</sup> | 6.6 (3.0)<sup>*</sup> | - |
| Superior Temporal Gyrus R GM (posterior parts) | 11.4 (2.4)<sup>***</sup> | 0.8 (0.3)<sup>*</sup> | - |
| Superior Temporal Gyrus L GM (posterior parts) | 10.8 (2.3)<sup>***</sup> | 0.8 (0.3)<sup>*</sup> | - |
| Anterior Cingulate Gyrus L GM | 6.4 (3.9) | -2.1 (0.83)<sup>*</sup> | -0.1 (0.0)<sup>*</sup> |

**Figure 4.**
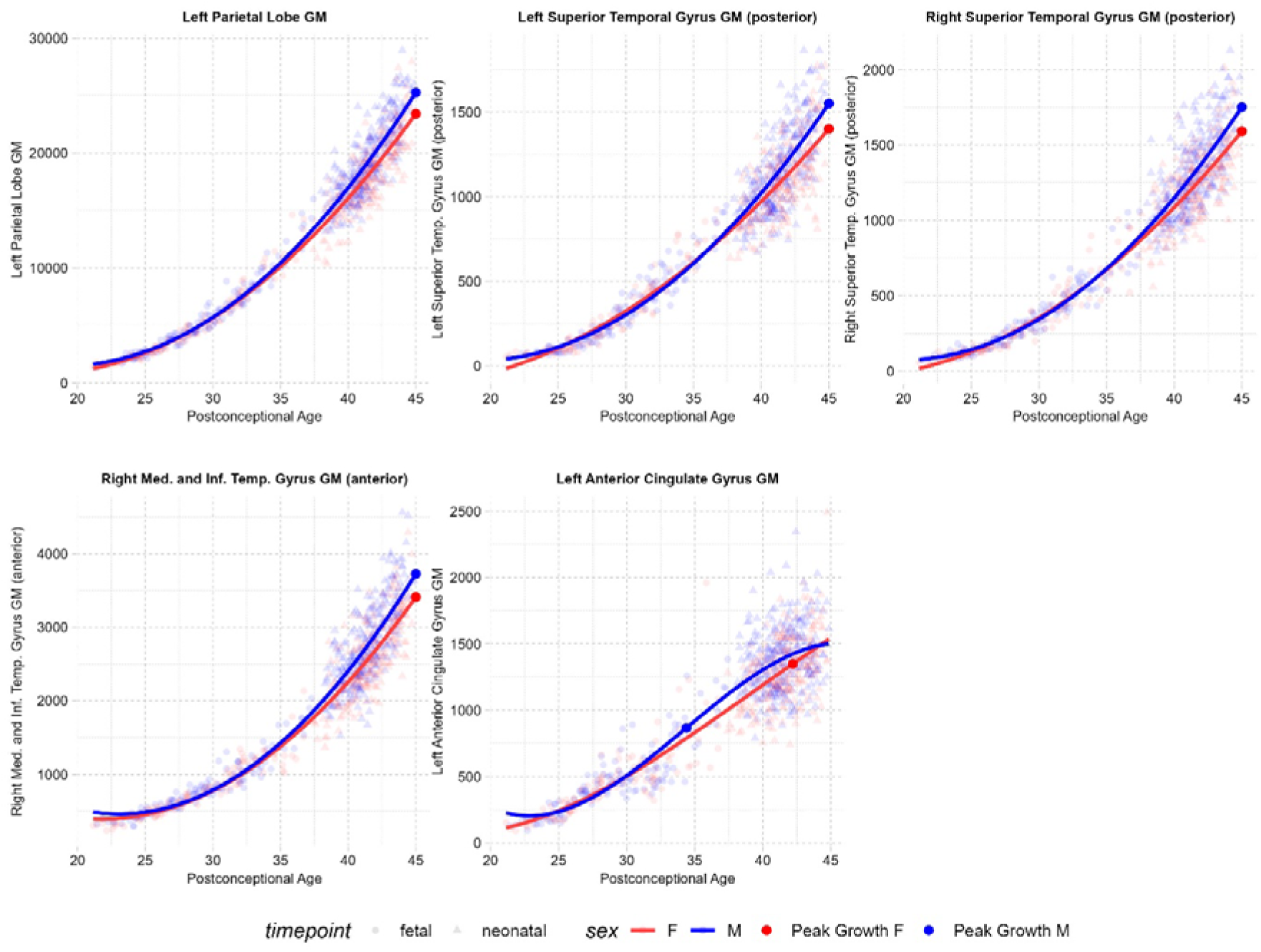
Regions Showing Significant Quadratic and Cubic Sex-by-Age Interactions.

Regional trajectories as a proportion of total brain volume are reported fully in Supplementary Materials (Supplementary Table 2). In summary, these trajectories followed the global trends reported above, with gray matter regional proportions increasing over time and white matter regional proportions decreasing over time. Various significant sex-by-age interactions were also evident, though in fewer regions compared to the absolute regional analysis.

## Discussion

The prenatal and early postnatal stages are among the most critical periods for brain development, yet these stages have seldom been studied continuously and in well-powered samples. In the present study, we modelled brain growth trajectories during the prenatal to postnatal transition. Using one of the largest prenatal and neonatal MRI datasets (798 scans), we identified various global and regional changes in volumetric dynamics between 21-45 weeks postconceptional age. The identified trajectories show that white matter dominates mid-gestational brain growth, while gray matter dominates late gestational and postnatal brain growth. Within gray matter growth, subcortical structures follow distinct trajectories and reach peak growth rates earlier than cortical structures. Additionally, the studied period captured a critical window for sex differences in brain growth, with males showing greater overall volumetric increases with age than females.

### Global Growth Patterns

The findings demonstrate that total brain volume increases at an accelerating rate between 21-45 weeks postconceptional age. A similar pattern is reported in prior prenatal studies, where brain growth is shown to rapidly increase between the late second trimester and term^9^. Postnatally, the rate of total brain growth is most rapid in the days immediately after birth and then slows down during subsequent weeks^7^. The second half of gestation and first few postnatal weeks are therefore particularly rapid windows for brain development, likely driven by processes such as dendritic and synaptic development, axonal outgrowth, and glial proliferation^1^. On the other hand, ICV and CSF both showed cubic trajectories. CSF levels were reduced when measured between 30-40 weeks and then increased at an accelerating rate from 40 weeks onwards. A transient drop in CSF volumes around birth has also been documented in prior studies^12^ and may be linked to the process of labour^28^. Given that the measure of ICV used here represented the sum of total brain volume and CSF, the observed cubic trajectory of ICV is likely also attributable to changes in CSF.

The findings also show that, between 21-45 weeks postconceptional age, total gray matter grows at an increasing rate while white matter grows at a decreasing rate. Notably, the relative contribution of these tissue classes to total brain volume changes across perinatal development. White matter is a greater contributor to total brain volume before ∼35 weeks, and proportional white matter overall declines across the studied period. Consistent with these findings, prior prenatal studies have shown that white matter is the greatest contributor to total brain growth during the second half of pregnancy, peaking between 29-30 weeks^9^. Moreover, diffusion imaging studies show that all major white matter tracts are present at birth, with certain tracts (e.g., thalamo-cortical fibres) already developed by the end of the second trimester^2,29,30^. White matter injury is also one of the most prominent brain-based differences between term-born and preterm infants^31^, reflecting the importance of prenatal white matter development. The subsequent decline in white matter growth rates has also been documented in prior research, where it is shown that white matter increases by only ∼11% from birth to 1 year, while gray matter increases by 108-149%^3,32^. Similarly, while gray matter volume reaches its peak in childhood, white matter volume only peaks in adulthood^33^. Rapid prenatal increases in white matter therefore likely establish core white matter connectivity, after which the myelination and maturation of these existing connections continue in a more gradual and protracted manner.

Proportional decreases in white matter coincide with proportional increases in gray matter. Gray matter is a greater contributor to total brain volume after ∼35 weeks, and proportional gray matter volumes overall increases across the studied age-period. These increases are likely due to rapid neural proliferation, dendritic spine arborization, neuropil maturation, and synaptogenesis^2^. These findings are also consistent with prior studies showing that gray matter volume increases more rapidly compared with white matter in the third trimester^9^ and postnatally^3,32,34^. Rapid gray matter growth during these periods may hold functional significance, likely enabling the development of key motor, sensory, and cognitive abilities to facilitate postnatal functioning.

### Absolute Regional Development

Following the global trends reported above, most cortical regions showed quadratic growth trajectories characterised by accelerating regional gray matter growth and decelerating regional white matter growth. Importantly, key differences were identified between regional cortical and subcortical gray matter development. While most cortical regions showed quadratic trajectories, subcortical regions such as the amygdala, thalamus, and basal ganglia structures showed cubic trajectories. These trajectories were characterised by an initial phase of accelerating growth during the third trimester, followed by a subsequent phase of decelerating growth postnatally. Moreover, these structures showed earlier peak growth rates compared with cortical gray matter structures. The relatively increased growth of these structures during the third trimester indicates that the prenatal environment plays a critical role in driving subcortical development, which, in turn, may be critical to orchestrating fundamental postnatal functioning. The importance of third trimester subcortical development is further emphasised by findings demonstrating that infants born very pre-term show reduced subcortical volumes at term-equivalent age, particularly in structures such as the thalamus and basal ganglia^35,36^. In turn, these reduced volumes have been associated with poorer cognitive, behavioural, and motor outcomes^36^.

Subcortical structures also include the cerebellum, which showed exponential growth throughout the studied age-period. The cerebellum contains more than half of the brain’s neurons at birth and is the region that shows the most rapid growth during the second half of gestation and early postnatal life^7,9^. Rapid perinatal cerebellar growth is likely integral to facilitating early motor coordination. Additionally, studies have shown that reduced early cerebellar volumes are associated with cognitive, motor, and socio-affective disruptions^37^, highlighting the cerebellum’s broader contribution to early functional development.

The hippocampus, on the other hand, was one of the few subcortical structures that showed a linear growth trajectory. Prior research has shown that the hippocampus is the slowest structure to mature during early postnatal development^7^. This is thought to reflect a more gradual development of processes such as memory formation and spatial navigation, which may be comparatively less critical for perinatal functioning (though remains crucial across longer-term development). Overall, these patterns suggest that perinatal regional growth mirrors early developmental patterns, with regions contributing to basic perinatal functioning (e.g., sensorimotor integration) showing earlier peak growth rates compared with regions contributing to higher-order cognition.

### Sex Differences

On average, males showed significantly faster prenatal and postnatal brain growth than females across the studied period. This pattern was reflected in all global brain volumes as well as several regional volumes. These findings are well-aligned with a large body of research that has shown that males have significantly larger brain volumes than females^16^, even as early as during the neonatal period^17^. Larger gray and white matter volumes in males have been associated with higher testosterone levels^38,39^. Similarly, research using an organoid model of the developing human brain has demonstrated that androgens induce cortical expansion^40^. The prenatal surge in testosterone in male fetuses peaks at around 14-18 weeks gestation and is followed by the first observable MRI-based sex differences in brain volumes at around 18 weeks gestation^41^. Observable sex differences in brain structure therefore appear a few weeks after these hormonal changes, likely due to the time required for intermediate cellular and molecular mechanisms to manifest into structural changes in brain volumes. Similarly, prior research has also measured prenatal anogenital distance (AGD), a proxy measure of androgen exposure, which is typically larger in males compared to females. Such research has demonstrated that sex differences in AGD begin prenatally and lag a few weeks behind the prenatal testosterone surge^42^.

Between 18-40 weeks gestation, the difference in testosterone levels in male and female fetuses remains consistent. Interestingly, a prior study which isolated the neonatal sample used in this research reported several main effects of sex on brain volumes at birth, but few sex-by-age interactions during the first month of postnatal life^17^. Since observable changes in brain volumes occur after hormonal changes, the limited sex-by-age interactions observed during the neonatal period may reflect the stabilisation of prenatal testosterone differences between males and females after mid-gestation. On the other hand, the inclusion of prenatal scans in the present study revealed several significant sex-by-age interactions, highlighting the prenatal period as a critical window for sex-differential growth.

Indeed, while sex differences in brain volumes are reported across the lifespan, sex-by-age interactions are most prominent only during a select few stages, particularly during puberty^43,44^ and prenatal development^18^. This is consistent with accounts suggesting that a few critical periods, often coinciding with sex steroid level changes, drive differential growth and produce sex differences that are subsequently observed throughout development^44,45^. Although males exhibit faster overall brain growth (captured by significant linear interactions), the shapes of the growth trajectories (captured by quadratic and cubic interactions) are largely similar between the sexes. In other words, both sexes generally follow similar global and regional trajectories as reported above, with the exception of a few regions. For instance, males exhibited more rapid and pronounced gray matter increases in the posterior parts of the bilateral superior temporal gyri and anterior parts of the medial and inferior temporal gyrus – regions which also show increased volumes in males at birth^17^. In the left anterior cingulate gyrus, males showed a cubic S-shaped trajectory with decelerating growth towards the end of the studied age-period, while females showed a linear trajectory with steady increases throughout. The region is consistently observed to be relatively larger in females across development^17,46^ and is typically associated with social cognition^47,48^, a domain in which females show advantages from early life^49^. It is important to note, however, that fewer sex-by-age interactions emerged when analysing regional volumes as a proportion of total brain volume, indicating that sex differences in absolute regional growth may be largely driven by the global sex difference in total brain volume.

It is possible that the growth patterns identified here shape sex-differential neurobiological development from early life, contributing to the emergence of sex differences in neurodevelopmental and psychiatric conditions later in life. While the exact mechanisms and causal pathways underlying this putative relationship remain insufficiently understood, sex hormones are thought play pivotal roles. For instance, fetal testosterone has been implicated in early-emerging conditions such as autism^24^, while the identification of several later-emerging conditions that show sex differences (e.g., depression, anxiety, eating disorders) also align with periods of significant hormonal changes, such as puberty^50^. This suggests that sex hormones are important modulators in the interplay between sex differences, brain development, and individual developmental outcomes. Analysing the developmental patterns of sex differences in the brain alongside the developmental patterns of neurodevelopmental and psychiatric conditions may help to explain the exact mechanisms driving these associations.

### Strengths, limitations, and future directions

There are important considerations that need to be taken into account when interpreting the findings of this study. First, the prenatal scans in the dataset begin after 21 weeks, limiting our understanding of brain growth during the first half of the prenatal period. Similarly, there were few prenatal scans after 37 weeks, limiting our understanding of brain growth in the final weeks of gestation in pregnancies lasting over 37 weeks. It is possible that brain development *in utero* after 37 weeks follows a somewhat different pattern to brain development *ex utero*, which may warrant further research. Second, although the dataset includes some longitudinal measurements, the longitudinal sample comprises only a subset of the full sample (∼14%). Further research on larger longitudinal samples will be important for validating the developmental trajectories identified here. Third, although both the prenatal and neonatal scans shared the same scanner and largely similar pre-processing procedures, their acquisition parameters differed. Commonly used harmonisation techniques (e.g., ComBat harmonisation) were deemed unsuitable in this case as they would likely not have been able to distinguish technical differences from true biological differences between prenatal and neonatal development. Nonetheless, we observed continuity across most of our growth curves, and the observed growth trajectories were consistent with those reported in prior prenatal and neonatal research, indicating that technical differences did not play a substantial role in the observed findings. Fourth, there were no available measures on fetal body size, and we were therefore unable to determine whether sex differences in prenatal brain growth are independent from sex differences in overall body growth. Nonetheless, it is noteworthy that prior studies on the neonatal period have reported that sex differences in brain size persist even after accounting for sex differences in birth weight^51^. Finally, the present research reports only one of many brain growth metrics. Further research incorporating additional neuroanatomical, diffusion-weighted, and functional measures will be important for gaining a more comprehensive understanding of perinatal brain development.

A considerable strength of the present research is the comparatively large sample size, making this one of the largest known studies on perinatal brain development. Importantly, this is also one of the few studies to analyse both prenatal and postnatal brain development at the same time, allowing us to map continuous perinatal growth trajectories. Additionally, while various existing studies on postnatal development begin from a few weeks after birth, the neonatal scans used in this research begin from day 0, capturing brain growth during the early neonatal period. The dHCP structural pre-processing pipeline^52^ used in this research is optimised for the studied period and overcomes several challenges typically encountered in prenatal and neonatal brain imaging (e.g., partial volume effects, low tissue contrast, motion artefacts). Importantly, both fetal and neonatal scans were pre-processed using the largely similar pipelines and segmentations, facilitating comparability across these life stages.

### Conclusion

In summary, using one of the largest prenatal and neonatal MRI datasets, our findings highlight changes in volumetric dynamics as the brain transitions from the prenatal to the postnatal stage. The identified patterns show earliest peak growth for white matter during the second trimester, followed by subcortical gray matter during the third trimester and, lastly, cortical gray matter postnatally.

Moreover, our findings indicate that shifts in volumetric dynamics predominantly occur during the final stages of gestation, suggesting that they are likely initiated by the prenatal environment in anticipation of birth. The studied age range also captured one of the critical periods for differential growth between males and females, with the timeline of the observed sex differences corresponding with the timeline of prenatal sex hormone fluctuations. Overall, the identification of these patterns emphasises the importance of integrating the prenatal and postnatal periods to facilitate a more comprehensive and continuous understanding of early growth trajectories.

## Methods

### Participants

Participants were recruited as part of the developing Human Connectome Project (dHCP)^25^, which was approved by the UK National Research Ethics Authority (14/LO/1169). In total, the dHCP dataset consists of 783 neonatal scans and 273 prenatal scans. Exclusion criteria employed for this study included multiple births, brain anomalies with likely analytical and clinical significance, any pregnancy or neonatal clinical complications, and preterm births (<37 weeks gestational age) for neonatal scans. The final sample used in the analysis consisted of 798 scans from 699 unique fetuses/neonates (380 males and 319 females, as assigned at birth) spanning a post-conceptional age range of 21.14 to 44.71 weeks. Amongst these were 263 fetal scans from 244 fetuses (131 males and 113 females) and 535 neonatal scans from 533 neonates (287 males and 246 females). Full sample characteristics are reported in Table 2. A subset of the sample (N=97) had longitudinal scans, of which 78 participants had both fetal and neonatal scans, 16 participants had 2 fetal scans, 1 participant had 3 fetal scans, and 2 participants had 2 neonatal scans.

**Table 2.** Sample Characteristics. *Sample characteristics for the full sample, also stratified by sex assigned at birth. P values for sex differences in sample characteristics, based on two-sample t-tests, are also reported. SD = standard deviation*.

|  | Range | Mean (SD)<br>All | Mean (SD)<br>Males | Mean (SD)<br>Females | Sex<br>differences *p*<br>values |
| --- | --- | --- | --- | --- | --- |
| Postconceptional<br>Age at Scan – all<br>scans | 21.14 – 44.71 | 37.43 (6.36) | 37.45 (6.25) | 37.46 (6.50) | 0.92 |
| Postconceptional<br>Age at Scan –<br>prenatal scans | 21.14 – 38.29 | 29.17 (3.90) | 29.24 (3.85) | 29.08 (3.97) | 0.72 |
| Postconceptional Age at Scan – postnatal scans | 37.43 – 44.86 | 41.50 (1.65) | 41.38 (1.60) | 41.62 (1.71) | 0.08 |
| Gestational age at birth – postnatal scans | 37 – 43 | 40.11 (1.18) | 40.07 (1.15) | 40.14 (1.22) | 0.48 |
| Postnatal Age at Scan – postnatal scans | 0 – 49 | 9.69 (9.44) | 9.15 (8.91) | 10.33 (10.00) | 0.15 |
| Birth weight – postnatal scans | 1.82 – 5.36 | 3.58 (0.55) | 3.65 (0.55) | 3.49 (0.55) | 0.002 |

### Data acquisition and preprocessing

#### Prenatal scans

All imaging data were acquired using a Philips Achieva 3T MRI scanner (Philips Medical Systems) with R3.2.2 software. Mothers were placed in a supine position, and their blood pressure and peripheral pulse oximetry was monitored throughout the scan. Structural T2-weighted (T2w) images were collected using a multi-band 2, single-shot Fast Spin Echo (ssFSE) sequence, which included an MB tip-back pulse to enhance signal-to-noise ratio (SNR). Data were collected from six uniquely oriented stacks centred on the fetal brain using the following parameters: Resolution = 1.5 x 1.5 x 4.0 mm (−1.1 gap), TR/TE = 2265+/250ms. Localised B1+ shimming was performed to optimise magnetic field uniformity, while B0 shimming was performed to correct for field sensitivity introduced by the tip-back preparation pulse.

3D T2w images were reconstructed with a 1.0 mm resolution from motion-affected ssFSE stacks using an automated slice-to-volume registration (SVR) pipeline. The processing pipeline included denoising, inhomogeneity correction using B1+ calibration, brain localisation via V-net, and multiple iterations of SVR and super-resolution reconstruction. Afterwards, the 3D images were reoriented to standard anatomical planes using a transformer network specifically designed for the fetal brain. Cubic interpolation was then applied to achieve a final resolution of 0.5 × 0.5 × 0.5 mm.

#### Neonatal scans

Imaging was performed on the same scanner using the dHCP neonatal brain imaging system, which featured a neonatal 32-channel phased array head coil and a custom-designed patient handling system from Rapid Biomedical GmbH (Rimpar, Germany). Infants were fed, swaddled, and then scanned without sedation. Earplugs (President Putty, Coltene Whaledent, Mahwah, NJ, USA) and neonatal earmuffs (MiniMuffs, Natus Medical Inc., San Carlos, CA, USA) were used for auditory protection. Heart rate, oxygen saturation, and temperature were monitored throughout the scan by a paediatrician or neonatal nurse As described in the dHCP protocol, a Cramer Rao Lower bound approach was used to maximise contrast to noise ratio. T2w inversion recovery Fast Spin Echo (FSE) sequences were acquired in sagittal and axial orientations. Relaxation times were set to 1800/150 ms for gray matter and 2500/250 ms for white matter^53^. The in-plane resolution was set to 0.8 × 0.8 mm², with a slice thickness of 1.6 mm and a slice overlap of 0.8 mm. Other parameters were as follows for T2w images: TR/TE = 12000/156 ms, SENSE factor 2.11 (axial) and 2.60 (sagittal).

The acquisitions were reconstructed using a motion correction algorithm, and transverse and sagittal images were fused into a single 3D volume for high resolution and accurate volume estimation^54^.

### Data preprocessing

Similar pre-processing procedures were used for both the prenatal and neonatal scans based on the dHCP structural pre-processing pipeline ^52^. In summary, the T2w images were motion-corrected, bias-corrected, and brain-extracted ^55^. Next, a probabilistic tissue atlas was registered to the corrected images. The Draw-EM algorithm ^56^ was then used for initial tissue segmentation into CSF, white matter, cortical gray matter, and subcortical gray matter. Subsequently, labelled atlases^57^ were registered to the images using both T2w images and gray matter probability maps from the initial segmentation. The final segmentation consisted of 87 gray and white matter structures^56–58^. Additional segmentation labels were available for the fetal scans, though these were not used in this analysis for the purposes of consistency with the neonatal segmentation. All fetal segmentations were also reviewed for anatomical accuracy, and labelling errors were manually corrected by a neuroscientist with expertise in fetal neuroanatomy. Preprocessing codes are all available online (https://github.com/BioMedIA/dhcp-structural-pipeline, https://github.com/MIRTK/DrawEM/tree/feature/fetal_segmentation).

### Analysis

Statistical analyses were conducted on R (version 4.3.3, 2024-02-29), using the packages *nlme, tidyverse*, and *ggplot2*. Mixed-effects models, with random intercepts modelled at the subject-level, were used to assess the effects of age, sex, and sex-by-age interactions on brain volumes. Additionally, in order to understand relative growth, regions and tissues were analysed as a proportion of total brain volume (regional analyses presented in supplementary materials). A Benjamini-Hochberg false discovery rate (FDR) correction was applied to each analysis category using a significance threshold of 0.05^27^. FDR corrections were applied separately for absolute and proportional analyses, as well as for global and regional analyses. A total of 87 cortical and subcortical gray and white matter structures were tested.

#### Model comparisons

Given the nonlinear nature of brain growth, polynomial models were considered to capture more complex growth trajectories. Prior to the analyses, Bayesian Information Criterion (BIC) values were computed for each brain measure (separately for absolute and proportional volumes) to determine whether a linear, quadratic, or cubic model best described its growth trajectory (Supplementary Tables 3 and 4). Compared to Akaike Information Criterion (AIC), BIC imposes greater penalties for model complexity to minimise overfitting and is recommended in scenarios where the primary aim is to select a model that best fits existing data as opposed to one that best predicts future observations^59^. Mixed-effects models were then fit to each brain measure using the model type with the lowest BIC value. To reduce multicollinearity in the polynomial models, postconceptional age at scan was mean-centred. The models were defined as follows:

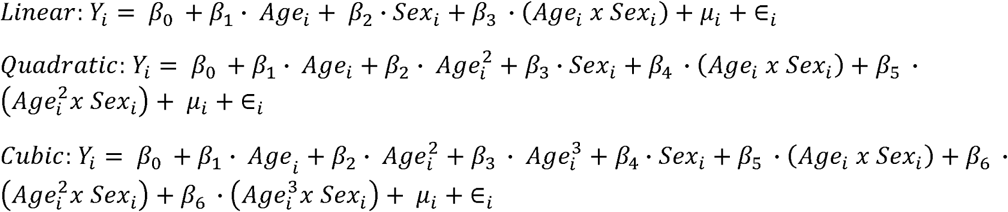

Where:

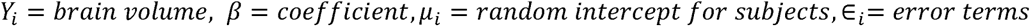

The same models were applied for proportional growth trajectories, where *Y_i_* represents the tissue or region as a proportion of total brain volume:

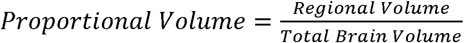

Standardised beta coefficients were also calculated for all analyses by z-scoring the data and are reported fully in the Supplementary Materials.

#### Variance modelling

As is well-documented in early developmental research^8,18^, the sample showed increasing variance in volumes with age. Visual inspection of the residual plots revealed a funnel-like shape, where the residual variance increased with age. To account for this, power variance weighting was applied using the *varPower* function in *nlme.* In this technique, each observation is weighted inversely proportional to a transformation of its variance, where the transformation is defined by a power function. This approach stabilises variance across ages, addressing heteroscedasticity and producing more reliable model estimates^60^.

#### Growth metrics

Peak growth points were calculated to identify the periods of most rapid growth within the studied age range. For quadratic models, peak growth points occurred at the extreme ends of the studied age range (beginning of the range for negative quadratic terms and end of the range for positive terms). For cubic models, peak growth points were determined via calculating the postconceptional age corresponding to the highest value of the first derivative.

## Acknowledgements

We thank Lena Dorfschmidt for providing a pre-processing script that aided the extraction of brain volume measurements for this analysis.

## Author contributions

Design and conceptualisation—All authors. Data processing: Y.T.K and A.T. Analysis: Y.T.K. Writing—original draft: Y.T.K. Writing—reviewing and editing: All authors.

## Materials & Correspondence

Yumnah T. Khan

